# HPV detection and genotyping of FFPE head and neck cancer biopsies by molecular testing to address new oropharyngeal squamous cell carcinoma classification based on HPV status

**DOI:** 10.1101/469387

**Authors:** David Veyer, Maxime Wack, Ophélie Grard, Pierre Bonfils, Stéphane Hans, Laurent Belec, Cécile Badoual, Hélène Péré

## Abstract

Recently, both the WHO/IARC (World Health Organisation/International Agency for Research on Cancer) and the American Joint Committee on Cancer (AJCC) have classified the oropharyngeal squamous cell carcinoma (OPSCC) on the basis of HPV status. For this purpose, the WHO/IARC recommended direct molecular HPV testing. In practice, formalin-fixed, paraffin-embedded (FFPE) biopsy specimens are frequently the only available samples. We herein compared in parallel two commercially available molecular assays that were firstly designed for cervical HPV detection and genotyping: Inno-Lipa^®^ HPV genotyping extra II assay (Fujirebio, Gent, Belgium) (IL) and Anyplex^TM^ II HPV 28 (Seegene, Seoul, South Korea) (AP28).

Both assays were carried out on the same DNA extracts obtained from prospectively collected FFPE biopsies from OPSCC origin and results were compared.

A total of 55 samples were tested. By IL assay, chosen as reference assay, 27 (49.1%) biopsies were positive for HPV16, 10 (18.2%) were positive for HPV but negative for HPV16, and 18 (32.7%) were negative for HPV. A valid result with AP28 was obtained for 51 biopsy samples (92.7%). Among 37 HPV-positive samples by IL, 33 (89.2%) were positive by AP28. The agreement between both assays was good (Cohen’s κ = 0.78). Among the six discrepancies between assays, always associated with low HPV16 viral load, four biopsies positive for HPV16 by IL could not be detected by AP28.

Taken together, these observations demonstrate that both assays could be used in routine for HPV detection and genotyping on FFPE-biopsy samples of head and neck tumour.

## Introduction

With 600,000 cases per year, head and neck cancer was estimated to be the sixth most common cancer worldwide (1). Head and neck squamous cell carcinoma (HNSCC) represents 90% of them. Since 2007, human papillomavirus (HPV) was considered as an independent risk factor for HNSCC by the World Health Organization’s International Agency for Research on Cancer (WHO/IARC) (2). Broad genetic distribution of HPV has been reported in oropharyngeal squamous cell carcinoma (OPSCC), including HPV16 in 85% of cases, followed by HPV18, 31, 33, 35, 45, 51, 52, 56, 58, 59, 68 and 82 (3, 4). Finally, OPSCC is currently considered as an epidemic viral-induced carcinoma, since the incidence of HPV-positive carcinoma of the tonsil nearly doubled every 10 years (5–8).

Recently, the major modification in the eight edition of the American Joint Committee on Cancer (AJCC) Staging Manuel, Head and Neck Section was the introduction of a specific staging algorithm for high-risk (HR) HPV-associated OPSCC (9). Furthermore, the 2017 edition of the WHO/IARC on Head and Neck tumor classified the OPSCC on the basis of HPV status (10–12). HPV-positive OPSCC constitutes a tumor entity with better prognosis, a distinct epidemiological profile, with specific genetic features and clinical presentations and outcomes. Immuno-histochemical detection of HPV using p16 staining as surrogate marker of HPV has been until now widely carried out (13). The 2017-revised WHO/IARC recommendations introduced direct HPV testing based on *in-situ* hybridization and/or PCR in order to classify the OPSCC according to HPV status (12). Until now, several commercially available assays have been clinically validated for the detection and genotyping of HPV-associated cervical cancer (14, 15), whereas to our knowledge none have been validated for OPSCC. Furthermore, formalin-fixed, paraffin-embedded (FFPE) biopsy samples are frequently the only available ones for molecular testing after pathological examination. Such FFPE samples however necessitate specific processing before PCR analysis because formalin fixation induces fragmentation of nucleic acids (16–18).

Taken together, the aim of the present study was to evaluate a new multiplex real-time PCR-based assay (Anyplex^TM^ II HPV 28, Seegene, Seoul, South Korea) (AP28), allowing to detect and genotype a wide range of high-risk HPV genotypes and previously tested for cervical samples (19, 20), in a prospective series of OPSCC FFPE samples, by reference to Inno-Lipa^®^ HPV genotyping extra II assay (Fujirebio, Gent, Belgium) (IL) chosen as reference assay (21–23).

## Materials and methods

### Collection of biopsy samples and processing

Head and neck biopsy samples received before treatment at the European Georges Pompidou hospital were prospectively included between 2014 and 2017 for routine pathologic examination. Patient had not received any treatment for their cancer at the time of the biopsy. The biopsies were fixed in formalin 10% overnight and included in paraffin. FFPE-biopsies diagnosed as HNSCC were selected by a pathologist for further cutting in 5- to 20- µm sections. Five sections were sent to the ISO 15189-accredited virology laboratory of the hospital for DNA extraction prior to HPV detection and genotyping by IL and, in parallel, by in-house quantitative real-time PCR targeting E6 gene from HPV16 (HPV16 qPCR). Afterwards, same DNA extracts were subjected to multiplex HPV PCR by AP28.

### DNA extraction procedures

Sections of FFPE-biopsies were deparaffinised overnight at +56°C with 40 µl of proteinase K (Qiagen, Hilden, Germany) and 360 µl of ATL buffer (Qiagen). Afterwards, 200 µl of ATL buffer were added and incubated 10 min at +70°C. DNA was further extracted using QiaAmp DNA Mini Kit (Qiagen), and eluted in 50 µl of PCR-grade water. For discordant results between IL and AP28, 5 new sections of biopsy samples were also subjected to DNA extraction procedure optimized for FFPE-biopsies, as previously described by Steinau *et al* (17).

### HPV detection and genotyping in routine

Two molecular HPV assays are used in parallel for routine HPV detection and genotyping on FFPE-biopsies. The IL assay consists of PCR amplification of a small 65 bp-fragment of the L1 gene using SPF10 primers sets and the ubiquitary gene human leukocyte antigen-DPB1 as internal control, followed by hybridization of specific HPV probes in a dedicated automat according to manufacturer’s instructions. The IL assay detects 13 HR HPV (HPV −16, −18, −31, −33, −35, −39, −45, −51, −52, −56, −58, −59, −68), 9 low-risk (LR) HPV (HPV −06, −11, −40, −42, −43, −44, −54, −61, −81), 7 genotypes reported as possibly carcinogenic (HPV −26, −53, −66, −67, −70, −73, −82) and 3 genotypes not described as carcinogenic (HPV −62, −83, −89) (24).

HPV16 qPCR was also systematically carried out, as previously described (25), in order to double check every sample for HPV16 which constitutes the most prevalent HPV genotype in HNSCC (26), as well as to assess lack of contamination by IL. For quantification, serial dilutions of titrated Caski cells (Amplirun, Orgentec, France) were used to plot external standard curve.

Positive controls for HPV16 and HPV18 consisted in DNA extracted from SiHa and HeLa cell lines, respectively; water was used as negative control.

### HPV detection and genotyping by multiplex PCR

The AP28 assay that distinguishes 28 HPV genotypes, by amplifying 100 to 200 bp-fragments of the L1 gene [including 13 HR types (HPV −16, −18, −31, −33, −35, −39, −45, −51, −52, −56, −58, −59, and −68), 8 LR types (HPV −6, −11, −40, −42, −43, −44, −54, −61) and 7 genotypes reported as possibly carcinogenic (HPV −26, 53, −66, −69, 70 −73 and −82)], and human gene β-globin in two different reactions was used for multiplex HPV molecular testing (27). Melting curves were obtained at 30, 40, and 50 cycles. Results were first automatically analysed using the Seegene Viewer software, version 2.0 (Seegene) and raw data of results were checked by the virologist. The results were considered as invalid when the negative controls were negative and no HPV was found. An estimation of the viral load was approached by indicating the cycle number at which the positivity was detected: + (50 cycles), ++ (40 cycles) and +++ (30 cycles).

### Ethical clearance

All included patients belonged to a cohort declared and approved by the Ethics Committee (Comité de Protection des Personnes Ile de France II, no. 2015-09-04)

### Statistical analysis

IL assay was chosen in our laboratory as the reference technique for HPV detection and genotyping, because the assay was reported to show high analytical sensitivity and specificity on FFPE samples (28, 29). Results strictly similar by AP28 and IL assays were defined as identical; results giving at least one identical HPV genotype by AP28 and IL assays were defined as compatible; other results were defined as discordant. Agreement between IL and AP28 assays was assessed by the Cohen’s κ test: 1 indicating perfect agreement; 1 to 0.81, very good agreement; 0.80 to 0.61, good agreement; 0.60 to 0.21; moderate to poor agreement. The Mann-Whitney test was used to compare HPV16 viral loads between samples showing concordant or discordant results. A linear regression model was used to assess the relation between the viral load and the semi-quantitative result of AP28.

## Results

Fifty-five biopsy samples from patients followed for HNSCC were prospectively selected. By the IL assay, 27 (49.1%) biopsies were HPV16-positive and 10 (18.2%) were HPV-positive but not HPV16-positive, as depicted in the Table 1. Among the 37 positive samples, 3 (biopsies #8, #19 and #33) were positive for more than one HPV. Finally, 18 (32.7%) biopsy samples were HPV-negative.

**Table 1.** HPV detection and genotyping by IL, AP28 and E6 HPV-16 detection and quantification by in-house PCR of 37 IL-positive FFPE-biopsies diagnosed as HNSCC. HPV 16-positive samples are presented by increasing viral load.

| ID | HPV genotyping results |  | In-house E6 HPV16 PCR |  |
| --- | --- | --- | --- | --- |
|  | IL | AP28<br>(semi-quantitative results<sup>a</sup>) | Qualitative results | Viral load<br>[copies/μl (log)] |
| #3 | HPV16 | Invalid<sup>b</sup> | Positive | ND |
| #5 | HPV16 | Invalid<sup>b</sup> | Positive | 133 (2.12) |
| #7 | HPV16 | Negative<sup>b</sup> | Positive | 503 (2.70) |
| #4 | HPV16 | Negative<sup>b</sup> | Positive | 512 (2.72) |
| #30 | HPV16 | HPV16 (+) | Positive | 734 (2.87) |
| #50 | HPV16 | HPV16 (++) | Positive | 3,910 (3.59) |
| #33 | HPV16 ; HPV82 | HPV16 (+) | Positive | 4,000 (3.60) |
| #9 | HPV16 | HPV16 (++) | Positive | 4,860 (3.69) |
| #14 | HPV16 | HPV16 (++) | Positive | 5,010 (3.70) |
| #46 | HPV16 | HPV16 (++) | Positive | 5,070 (3.70) |
| #10 | HPV16 | HPV16 (++) | Positive | 5,360 (3.73) |
| #45 | HPV16 | HPV16 (++) | Positive | 5,650 (3.75) |
| #35 | HPV16 | HPV16 (++) | Positive | 7,730 (3.89) |
| #13 | HPV16 | HPV16 (+) | Positive | 9,230 (3.97) |
| #49 | HPV16 | HPV16 (++) | Positive | 20,200 (4.31) |
| #44 | HPV16 | HPV16 (++) | Positive | 20,800 (4.32) |
| #12 | HPV16 | HPV16 (+) | Positive | 35,200 (4.55) |
| #2 | HPV16 | HPV16 (+) | Positive | 40,200 (4.60) |
| #48 | HPV16 | HPV16 (++) | Positive | 78,300 (4.89) |
| #11 | HPV16 | HPV16 (++) | Positive | 78,400 (4.89) |
| #32 | HPV16 | HPV16 (++) | Positive | 80,200 (4.90) |
| #16 | HPV16 | HPV16 (++) | Positive | 82,400 (4.92) |
| #6 | HPV16 | HPV16 (++) | Positive | 112,000 (5.05) |
| #27 | HPV16 | HPV16 (++) | Positive | 128,000 (5.11) |
| #39 | HPV16 | HPV16 (++) | Positive | 183,000 (5.26) |
| #1 | HPV16 | HPV16 (++) | Positive | 323,000 (5.51) |
| #42 | HPV16 | HPV16 (+++) | Positive | 595,000 (5.77) |
| #34 | HPV6 | HPV6 (++) | Negative | NA |
| #43 | HPV6 | HPV6 (++) | Negative | NA |
| #18 | HPV11 | HPV11 (++) | Negative | NA |
| #31 | HPV11 | HPV11 (++) | Negative | NA |
| #40 | HPV11 | HPV11 (++) | Negative | NA |
| #19 | HPV18 ; HPV39p | HPV18 (++) | Negative | NA |
| #47 | HPV18 | HPV18 (+++) | Negative | NA |
| #8 | HPV33 ; HPV52p | HPV33 (+++) | Negative | NA |
| #38 | HPV59 | HPV59 (++) | Negative | NA |
| #22 | HPV82 | HPV82 (+++) | Negative | NA |

<sup>a</sup> Semi-quantitative results are given in brackets (detection of signal at 30 cycles: +++ ; 40 cycles: ++ ; 50 cycles: +);
<sup>b</sup> Results checked manually without the automated analysis software (Seegene Viewer version 2.0, Seegene).
IL: Inno-Lipa® HPV genotyping extra II assay (Fujirebio, Gent, Belgium); AP28: Anyplex™ II HPV 28, Seegene (Seoul, South Korea); NA: Not attributable; ND: Not done; p: probable

### Comparison of HPV detection and genotyping results between IL and AP28 assays

The same DNA extracts from the 55 selected biopsy samples tested by IL were further subjected to AP28 assay. Among 7 discordant samples, one that was initially negative when analyzed with the Seegene Viewer software was finally classified as positive after raw data analysis; the raw data analysis of the remaining discordant samples (corresponding to 10.9% of samples) gave similar results to the ones obtained by automatic analysis. Final results are shown in the Tables 1 and 2. A valid result with AP28 was obtained for 51 biopsy samples (92.7%) (Table 2). Among the 37 HPV-positive biopsy samples by IL, 33 (89.2%) were found positive by AP28, including 90.9% of identical results and 9.1% of compatible results, the remaining results being either invalid (n=2) or negative (n=2). Finally, the vast majority (n=16; 88.9%) of 18 biopsy samples negative by IL were also negative by AP28; only 2 IL-negative biopsies were found invalid by AP28. The overall agreement between both assays was good (Cohen’s κ coefficient = 0.78). Among the 27 samples that were HPV16-positive with IL, AP28 detected HPV16 in 23 (85.2%), 2 (7.4%) were negative and 2 (7.4%) invalid (Table 1). AP28 did not detect any HPV16 in samples that were negative for HPV16 with IL. Regarding HPV16 detection, the agreement between both assays was good (Cohen’s κκcoefficient=0.75).

**Table 2.** Qualitative detection of HPV by IL and AP28 asays in 55 FFPE-biopsies diagnosed as HNSCC.

| IL/AP28 results | N (%) |
| --- | --- |
| **Concordant results** |  |
| . Positive/Positive | 33 (60.0) |
| ○ Identical | 30 (90.9)<sup>a,b</sup> |
| ○ Compatible | 3 (9.1)<sup>c</sup> |
| . Negative/Negative | 16 (29.1) |
| **Discordant results** |  |
| . Positive/Negative | 2 (3.6) |
| . Positive/Invalid | 2 (3.6) |
| . Negative/Invalid | 2 (3.6) |
| **Total** | **55 (100.0)** |

<sup>a</sup> In brackets: percentage of IL/AP28 concordant out of total Positive/Positive;
<sup>b</sup> Among discordant samples, only one initially negative by Seegene Viewer software (Seegene) showed raw data compatible with positivity, and was classified as positive sample;
<sup>c</sup> In brackets: percentage of IL/AP28 compatible out of total Positive/Positive.
IL: Inno-Lipa® HPV genotyping extra II assay (Fujirebio, Gent, Belgium); AP28: Anyplex™ II HPV 28, Seegene (Seoul, South Korea)

### HPV16 viral load

HPV16 qPCR was carried out on the 55 biopsy samples. All HPV16-positive samples with IL were positive by HPV16 qPCR and all HPV16-negative samples with IL were also negative with HPV16 qPCR (Table 1). The median of HPV16 viral loads was higher in concordant than in discordant samples [20,800 copies/µl (4.32 log/l), range, 734-595000 *versus* 503 copies/µl (2.70 log/l), range, 133-512; P=0.00077]. Interestingly, the HPV16 viral loads of IL-positive/AP28-negative results were low (Table 1). As expected, using a linear regression model, a significant relation (P=0.000388) was observed between the viral load (expressed in log_10_) and the AP28 semi-quantitative results with a linear coefficient of 0.79 log between each level of AP28 (negative, +, ++ and +++). The Figure 1 depicts the results of HPV16 viral loads and AP28 semi-quantitative results.

**Figure 1.**
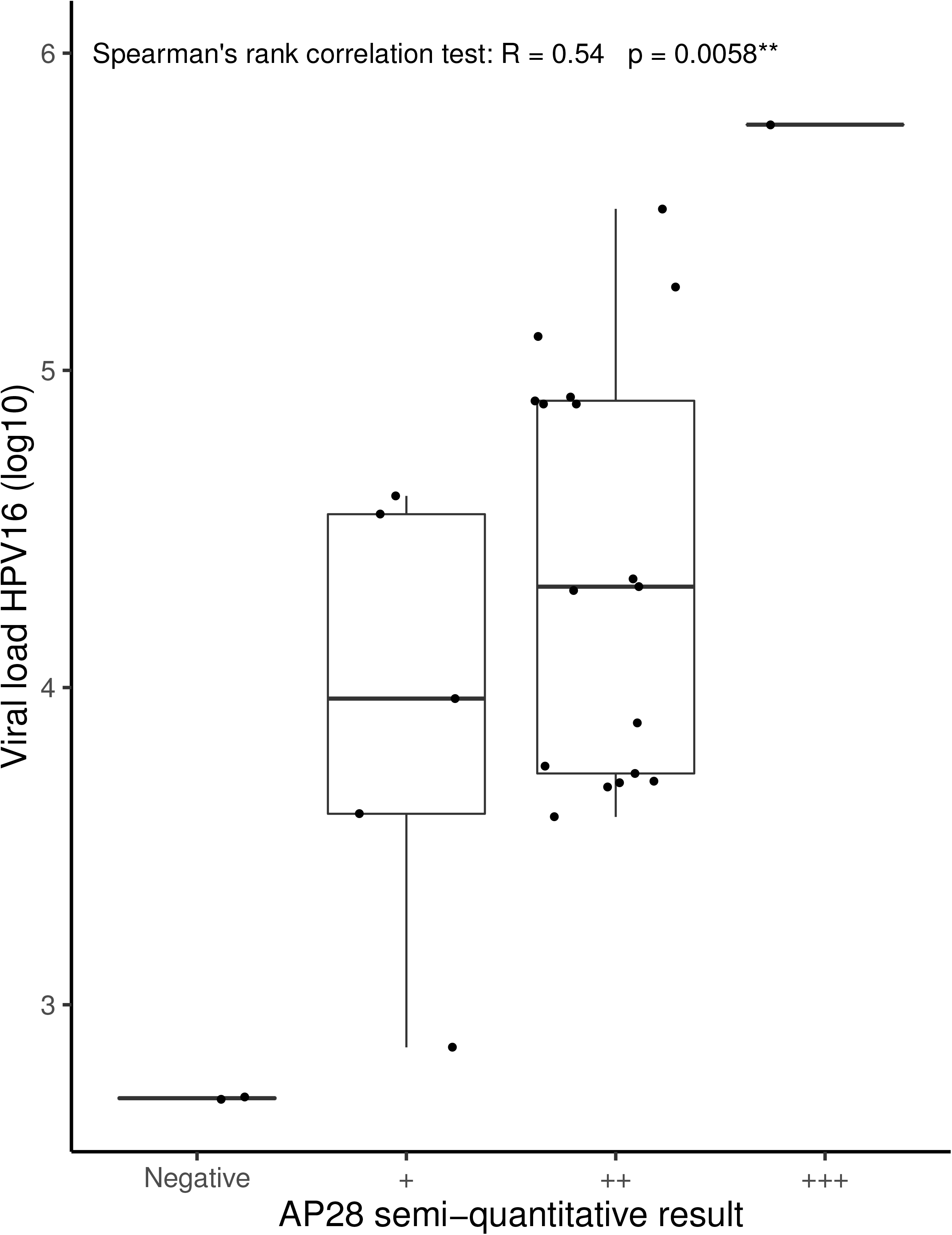
Comparison of HPV16 viral loads (in-house qPCR) and HPV16 AP28 semi-quantitative results. HPV16 semi-quantitative results are presented as negative, +, ++ and +++ according to the cycle at which the signal is detected (no detection, detection at 50, 40 and 30 cycles respectively). The Spearman’s rank correlation test is shown on the figure (R=0.54, p=0.0058).

## Discussion

We herein compared on a large series of FFPE-biopsy samples of HNSCC two commercially available molecular assays targeting L1 gene for routine HPV detection and genotyping. Nearly half of FFPE-biopsy samples from patients followed for HNSCC in our hospital was found positive for HPV DNA using IL as reference assay, emphasizing the need to strictly follow the recent 2017-revised WHO/IARC recommendation of direct HPV testing in case of OPSCC. Both assays showed good agreement for HPV detection as well as for genotyping. However, rare discrepancies between assays could be observed with positive samples for HPV16 by IL which were not detected by AP28. Interestingly, these discrepancies were always associated with low HPV16 viral load. These observations demonstrate that both assays could be used in routine for HPV detection and genotyping on FFPE-biopsy samples of HNSCC, keeping in mind that a few biopsy samples with low HPV16 viral load could be missed. Taken together, our findings clearly emphasize the necessity to validate commercially available molecular assays on patients’ biopsy samples, and to confront the results with other molecular technics in case of low HPV viral load.

In the present series, the calculated agreement between IL and AP28 results was good, both for HPV16 and other genotypes. However, around ten percent of biopsy samples gave discrepancies between both assays, including biopsies positive for HPV16 by IL which were not detected by AP28. Using HPV16 qPCR, low HPV16 viral load was clearly associated with the misdetection of HPV16 by AP28 assay. Although HPV16 viral load was not normalized on extracted DNA quantity, the experimentations were carried out on the same tissue extract allowing accurate comparison of the results obtained by AP28 and HPV16 qPCR. The capability of IL assay to detect HPV16 in samples harboring low HPV16 viral load is likely due to sufficient amplification efficiency in samples containing fragmented DNA. Indeed, it is well reported that DNA recovering in FFPE specimens may be influenced by several factors, such as formalin quality and concentration, length of fixation, paraffin quality and temperature (30). As a consequence, DNA in FFPE biopsy is either completely or partially degraded into DNA fragments of 200 bp or less (16). The IL assay relies on the amplification of shorter fragments than AP28 assay, a feature which could partly explain the discrepancies observed between both assays in our series. Furthermore, discordant results could also be associated with extract quantity used for experiment. Thus, the IL assay requires more quantity of tissue extract (10 µl) than AP28 (5 µl), according to manufacturers’ instructions. The possibility may be also envisioned that mutations affecting the priming sites of AP28 primers could have led to misdetection. This hypothesis should be however ruled out because the correlation between the efficiency of detection by AP28 and HPV16 viral load was marked. Finally, we checked that observed discrepancies were not due to extraction procedures.

In our series, one negative sample by AP28 which was positive by IL was finally diagnosed as positive for HPV16 after analysis of the raw data. This finding indicates that raw data of every AP28 negative samples need to be analyzed in order to detect some of the false-negative results. Indeed, the automatic analysis and cut-off used by AP28 test may miss out some HPV-positive samples since the assay was primarily designed to detect HPV on either cervical swab or liquid based cytology specimen and not on FFPE samples from oropharyngeal origin. In practice, the automatic Seegene software interpretation should be used with caution for negative or invalid FFPE biopsies.

Our observations indicate that IL as well as AP28 assays could be used in clinical laboratory to detect and genotype HPV in FFPE OPSCC biopsy samples. To our knowledge, only one study compared the performance of the AP28 assay and the CLART system from Genomica (Madrid, Spain) on only 3 FFPE HNSCC samples (27). In our series, the IL assay appeared as sensitive as in-house HPV16 qPCR, in keeping with previous reports showing that IL is highly reliable on FFPE archival samples. Nevertheless, the AP28 technic could be preferred depending on the recruitment of the laboratory, particularly if the number of samples is important. Indeed, AP28 relies on real-time PCR, which allows easier and faster procedure as compared with IL technic which appears more cumbersome and requires specific material for hybridization on membrane following PCR amplification. In addition, the risk of cross contamination between samples is less important with AP28 than with IL. However, it should be kept in mind that misdetection of HPV16-positive biopsies could occur with AP28 assay. Since HPV16 is known to be the most prevalent HPV genotype in OPSCC, diagnosed in 85% of cases (26), one could strongly recommend confirming the negative results by AP28 with HPV16 genotype-specific molecular assay validated on FFPE samples. In our laboratory, we have chosen to screen first for HPV16 by in-house HPV16 qPCR, and then to test further the negative samples with IL or AP28 to potentially detect less common genotypes in OPSCC, or confirm the negativity for HPV.

In conclusion, our observations demonstrate that both commercially available IL and AP28 assays could be used in clinical laboratories with similar performances in routine on OPSCC FFPE samples, in order to address the recent recommendations for direct HPV testing in OPSCC (12). Nevertheless, low HPV viral load in FFPE biopsy samples could be a limiting factor, rendering in medical practice the diagnosis of HPV in OPSCC sometimes difficult. Finally, HPV diagnosis and genotyping in OPSCC may necessitate several complementary technics, to overcome the limitation of some commercially available assays.

## Authors’ contributions

DV, OG, CB and HP have conceived and designed the research; OG, has performed the experiments; MW performed statistical analyses; DV, PB, SH, CB and HP analyzed the results; DV, LB and HP drafted the manuscript.

